# Coupled functional physiological phenotyping and simulation model to estimate dynamic water use efficiency and infer transpiration sensitivity traits

**DOI:** 10.1101/2022.10.10.511465

**Authors:** Ting Sun, Rui Cheng, Yudong Sun, Rujia Jiang, Zhuoyi Wang, Pingping Fang, Xinyang Wu, Kang Ning, Pei Xu

## Abstract

As agricultural drought becomes more frequent worldwide, it is essential to improve crop productivity whilst reducing the water consumption to achieve a sustainable production. Plant transpiration rate and water use efficiency (WUE) collectively determine the yield performance, yet it is challenging to balance the two in breeding programs due to still insufficient mechanistic understanding of the traits. Here we demonstrate the feasibility and effectiveness of calculating dynamic and momentary WUE by coupling WUE model and the state-of-the-art functional physiological phenotyping (FPP). We also present the method of quantifying genotype-specific traits reflecting sensitivity of transpiration to radiation (S_Tr-Rad_) and vapor pressure deficit (S_Tr-VPD_), under evolving developmental stage and water availability. Using these methods, we revealed the genotypic difference of S_Tr-Rad_ and S_Tr-VPD_ in three watermelon accessions, the dramatic change in each of them across the drought treatment phases, and the quantitative impacts of them on dynamic WUE patterns. Based on our results and computational simulations, a general principle for transpiration ideotype design is proposed, which highlights the benefits of lowering S_Tr-VPD_ to increase WUE and increasing S_Tr-Rad_ to offset the decline of Tr. FPP-enabled phenomic selection will help screen for elite crops lines with desired transpiration sensitivities.

## 1. Introduction

Global farming land area subject to drought is increasing due to the enhanced evapotranspiration associated with increased temperature, net radiation and decreased relative humidity (IPCC, 2021). Improving crop productivity, particularly under deficit water condition, is essential to safeguard food security under the context of global climate change. Stomata is the gateway of both water efflux (transpiration) and CO_2_ intake (assimilation). The plant transpiration rate (Tr) and the ratio of CO_2_ assimilation to Tr, which is known as water use efficiency (WUE), collectively determine biomass production and yield (Steduto *et al*., 2012; Vadez *et al*., 2014). Large efforts are being devoted to developing drought-resistant crops by adopting a strategy of restricting stomatal closure and thereby water loss; however, a tradeoff between increased WUE and decreased total transpiration as a consequence of stomatal restraint has long impeded breeding for agronomically useful varieties (Dalal *et al*., 2019). Previous studies have stressed that high-WUE crops usually had a smaller leaf area index (LAI), shorter development stage and thus lower Tr and carbon assimilation (Blum, 2005; Blum, 2009).

Traditionally, WUE is considered to be a conservative genotype-dependent trait (Steduto *et al*., 2012) and is measured by carbon-isotope discrimination (Moghaddam *et al*., 2013) or canopy gas exchange measurement systems (Song *et al*., 2016). It is now increasingly recognized that WUE is highly variable in response to environment change, developmental stages, and abiotic stress (Nelson *et al*., 2018; Peddinti *et al*., 2019; Yang *et al*., 2021). Monitoring dynamic WUE is therefore becoming routine in physiological studies and crop cultivation practices, even though it is presently very challenging due to technical constraints in the continuous nondestructive measurement of carbon assimilation and transpiration processes. WUE can change with the pattern of Tr within a day even if the daily cumulative Tr remains unchanged (Taylor *et al*., 1983; Vadez *et al*., 2014; Ghanem *et al*., 2020). Sinclair and Vadez (2012) decoded daily WUE as a function of vapor pressure deficit (VPD) weighted by diurnal Tr, implying the feasibility of selecting high-WUE varieties by harnessing genotypic variations in the dynamic Tr profile. It has been confirmed that restricted maximum Tr at noon under high VPD could increase the WUE and yield of maize (Jafarikouhini *et al*., 2022), sorghum (Sinclair *et al*., 2005) and soybean (Sinclair *et al*., 2010).

Mechanistically, Tr is not only governed by internal factors such as LAI and structure (e.g. cuticle thickness, stomata number), but external factors including temperature, relative humidity, light intensity, wind speed and soil water availability (Kubota, 2020). The interaction between Tr and the environment confounds phenotypic selection for Tr. According to Pieruschka *et al*. (2010), transpiration is mainly controlled by the diffusion gradient of water vapor inside and outside the stoma driven by VPD, and internal evaporation driven by latent heat from Rad. Since the diurnal patterns of Rad and VPD are inconsistent (Gosa *et al*., 2019), the sensitivities of Tr to VPD and Rad would have a strong impact on the dynamic Tr profile. Simulation models, which combine many biological functional hypotheses in mathematical frameworks, describe the development of plant traits as a consequence of environmental and genetic interaction (Génard *et al*., 2016). Several simulation models for Tr have been established (Katsoulas & Stanghellini, 2019). Recently, simulation models have found their new use in inferring parameters representing genotype-specific traits or “hidden traits” that are not intuitively extractable from the measured phenotypes. Such an approach is referred to as “inverse modeling” (Chenu *et al*., 2017).

Compared to morphological traits, plant physiological traits are notoriously difficult to phenotype. The emerging functional physiological phenotyping (FPP) has offered revolutionary paths for phenotyping plant water relations (Xu *et al*., 2015; Halperin *et al*., 2017; Chenu *et al*., 2018; Li *et al*., 2020). One of such platforms named “Plantarray” allows for high-resolution (every 3 minutes), non-destructive, simultaneous and continuous recording of meteorological variables and physiological traits such as diurnal Tr and daily biomass gain based on the lysimeter principle. The acquired massive dynamic data from the soil-plant-atmosphere continuum provides unprecedent opportunities for deeply understand the mechanisms of plant water relations. Here, we report on the estimation of dynamic WUE, a traditionally difficult-to-measure trait, by coupling existing WUE models with FPP, as well as the quantification of the genotype-specific dynamic traits of “Tr sensitivity to Rad/VPD” (S_Tr-Rad_ and S_Tr-VPD)_ by coupling FPP and the modified Penman–Monteith model (Medrano *et al*., 2005; Jo & Shin, 2021). This study demonstrates the great promise of leveraging advanced phenotyping techniques and existing modeling methods to tackle the long-standing complex trait challenge. The implications for transpiration ideotype design are discussed.

## 2. Materials and methods

### 2.1. Plant materials and experimental design

Three genotypes of watermelon, i.e., a wild watermelon accession “PI-296341-FR” (*Citrullus lanatus* var. citroides), a cultivated watermelon variety “Jincheng No. 5” (*Citrullus lanatus* var. lanatus, hereafter “JC5”) popular in Northwest China, and a drought-tolerant inbred line of *Citrullus lanatus* var. lanatus “HA” were used in this study. The experiments were carried out in the spring and summer seasons of 2021 in a semi-controlled greenhouse (L×W×H: 10m×5m×4m) in Huai’an (119°01′E, 33°35′N), China. One seedling at the three-leaf stage was transplanted per pot (16cm×13cm×18cm, 1.5 L) with Profile Porous Ceramic substrate (diameter: 0.2mm, pH: 5.5±1, porosity: 74, and CEC: 33.6mEq/100g).

Four replicates for each genotype were set in a completely randomized design on “Plantarray” (Plant-Ditech, Israel, Fig.1a). This system provides a simultaneous measurement and detailed physiological response profile for plants in each pot (Dalal *et al*., 2020). Water and nutrient supplements (Yamazaki nutrient solution) were controlled by the automatic irrigation system of Plantarray. Experiment in each growth season included three treatment periods, i.e., well-irrigation (WI), progressive water deficit (WD) and water recovery (WR). Under the WI and WR phase, nutrient solution was provided by irrigation for 240 seconds (oversaturated) at 23:00, 1:00, 2:00, and 3:00 of a day, respectively, and no nutrient solution was supplied during the WD stage.

### 2.2. Functional physiological phenotyping (FPP)

The Plantarray system is a weighing type lysimeter that consists of highly sensitive, temperature compensated load cells (Halperin *et al*., 2017). A series of functional physiological traits relative to the water loss by plant transpiration and plant growth could be obtained by the weight change throughout the day using the SPAC software implemented in the Plantarray system (Fig. 1a & b). Briefly, the pre-dawn system weight (averaged over 4:00-4:30 a.m.) was recoded as W_m_ at the end of the free drainage after saturated irrigation, and system weight in the evening (averaged over 9:00-9:30 p.m.) was recoded as W_e_ before irrigation, both of which were stable due to no water loss by plant transpiration and water input from irrigation during this period. As each pot on the system was covered with plastic film, soil evapotranspiration was prevented. The whole-plant daily transpiration (*E*, g d^-1^) was calculated as the difference between the W_m_ and W_e_ for each day. The daily plant growth (*PG*, g FW d^-1^) was determined as the difference of W_m_ between consecutive days, as the soil weight at field capacity in different days remained constant. The whole-plant WUE during the well irrigation period was thus determined by the ratio between the accumulated *PG* and *E*. The momentary whole-plant transpiration (Tr, g min^-1^) at 3-min step was calculated by multiplying the first derivative of the measured load-cell time series by -1, assuming that the plant’s weight gain during the short time interval used to calculate the transpiration rate was negligible (Halperin *et al*., 2017).

**Figure 1.**
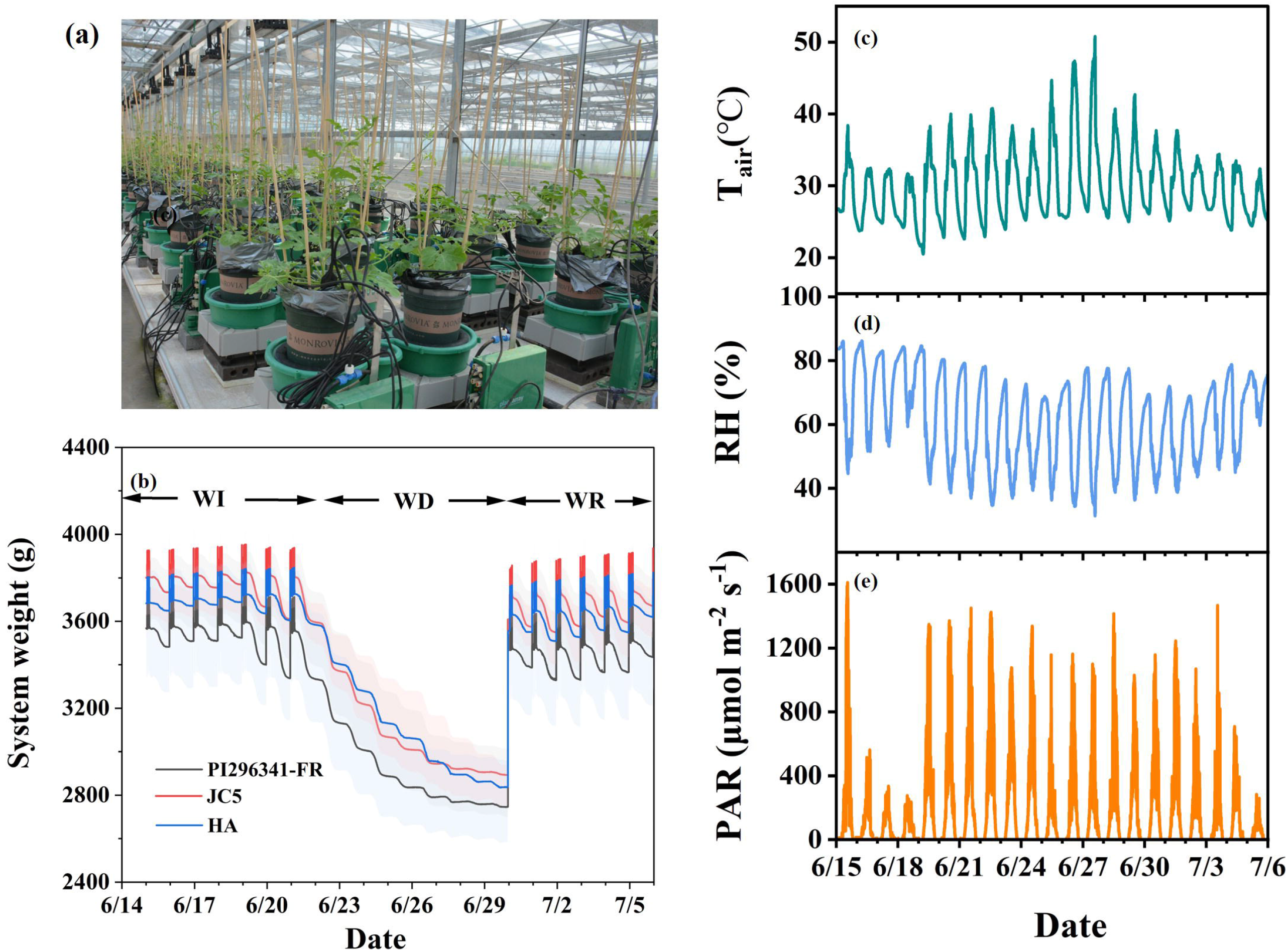
Parameters of soil–plant–atmosphere continuum monitored by high-throughput phenotyping platform “Plantarray”. a, overview of the automated physiological phenotyping array loaded with watermelon seedlings. b, system weights that were recorded every 3 minutes during the experimental course. Shadow areas show the observed variation. The experiment was divided into the well-irrigation (WI), progressive water deficit (WD), and water recovery (WR) phases. c-e, meteorological parameters, i.e., air temperature (T_air_, L, c), relative humidity (RH, %, d), and photosynthetic active radiation (PAR, μmol m^-2^ s^-1^, e) during the summer-season experiment in 2021.

The air temperature (T_air_, □) and relative humidity (RH, %), photosynthetically active radiation (PAR, μmol m^−2^ s^−1^) above the canopy, and soil volumetric water content (VWC, m^3^ m^−3^) in each pot were measured by the VP-4, 5TM, and PYR solar radiation sensor (Decagon Devices, Pullman, Wash, USA), respectively (Fig. 1c-e, Fig. S1). VPD (kPa) was calculated according to Halperin *et al*. (2017).

### 2.3. Simulation of WUE and biomass gain

The initial definition of WUE considers the resistances to diffusion resulting from stomata, leaf aerodynamic boundary layer, and leaf mesophyll resistance, and can be ultimately simplified to:

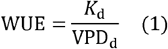

where VPD_d_ is the mean VPD over a day. *K*_d_ is a coefficient that reflects the CO_2_ concentration in the stomatal chamber, which has been validated among different crops (Tanner & Sinclair, 1983; Sinclair *et al*., 1984). *K*_d_ was estimated as 6.5 kPa in this study for watermelon plant with the data of FPP and WUE obtained from destructive sampling during a long growth period.

To estimate WUE over a long period, VPD needs to be integrated over a growth season and weighted by the transpiration over the course (Taylor *et al*., 1983; Vadez *et al*., 2014; Ghanem *et al*., 2020). Applying the same logic to a single day, Sinclair and Vadez (2012) proposed an assumption that the weighting of this component based on the transpiration rate throughout the day leads to the possibility of obtaining altered values if there are genotypic variations in the transpiration profile throughout the day. Therefore, during a computation interval (i), the mean VPD (VPD_i_) is weighted for mean transpiration rate (Tr_i_) to calculate the estimated WUE (WUE_e_) as in the following integrating equation:

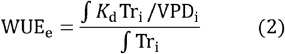

And biomass gain during this period could be estimated as:

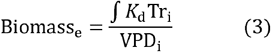

### 2.4. Canopy gas exchange parameters for validation

Gas exchange parameters were obtained by the classical but low-throughput canopy photosynthesis and transpiration system (CAPTS 100, China) from wheat plants which were cultivated in the field in 2021. The gas exchange parameters of the 1 m × 1 m canopies were measured automatically every 30 min using CAPTS 100 consisting of a cubic transparent chamber, which can be open and closed automatically with programming and equipped with CO_2_, humidity, air pressure, and temperature sensors in it (Song *et al*., 2016). Canopy gas exchange rate was measured according to the change rate of CO_2_ and H_2_O concentrations in the chamber during the closure of the chamber (Fig. S2a). The parameter *K*_d_ for wheat plants in this study was estimated using the least square method by fitting the estimated and measured daily WUE obtained during the growth period of March; then estimated and measured WUE at half-hour intervals and daily intervals during the days between April 5 and April 10 of 2021 were shown in Fig. S2b.

### 2.5. Transpiration model

We choose the simulation models which account for the response of Tr to the major meteorological variables and plant growth, following the principle that model obtains simple structure using as few parameters as possible. A modified Penman–Monteith model which is appropriate for greenhouse environment, is employed to simulate the Tr of watermelons in this study, which has been test for cucumber and tomato plants (Medrano *et al*., 2005; Jo & Shin, 2021). The diurnal Tr could be described as a function of radiation, VPD and leaf area index (LAI):

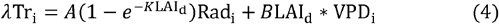

where λ is the vaporization heat of water (J kg^-1^), Tr_i_ is the instantaneous transpiration rate, *K* is the light extinction coefficient (0.86), Rad_i_ is the instantaneous solar radiation (W m^-2^, converted from PAR), LAI_d_ is the daily leaf area index (m^-2^ m^-2^). VPD_i_ is the instantaneous vapor pressure deficit. The LAI was estimated based on daily transpiration in each experiment unit by considering the individual difference (detailed as follows).

### 2.6. LAI estimation and validation

Step 1, dynamic Tr recorded in a frequency of every 3 minutes across a day was fitted to the observed radiation and VPD, thus optimum *A, B*, and LAI in the modified Penman–Monteith equation (Eq. 4) were estimated according to nonlinear least square (NLS) method and trust-region algorithm via the trial version of MATLAB 2018b (MathWorks, Inc., Cambridge, MA, USA). Only daily LAI with robust goodness of fit (adjusted R^2^>0.85) were picked for subsequent calculations. Step 2, we checked the accumulated daily transpiration (*E*) measured by Plantarray and the estimated LAI from daily fitting of step 1, where good linear relationship was found between them. Therefore, a constant LAI growth rate per E was estimated for individual experiment units, and then daily LAIs of individual experiment units were re-calculated according to daily measured E and estimated LAI growth rate per E. Step 3, daily parameter *A* and *B* were determined with observed Tr, RAD, VPD and re-calculated LAI during Step 2 for WI, WS, and WR stage, respectively. Parameter fitting was performed also with MATLAB 2018b.

Independent experiment was conducted to verify the aforementioned method, where estimated LAI was compared to the LAI measured by an LA-S leaf area analyzer system (Wanshen Technology Co., Hangzhou, China, Fig. 2).

**Figure 2.**
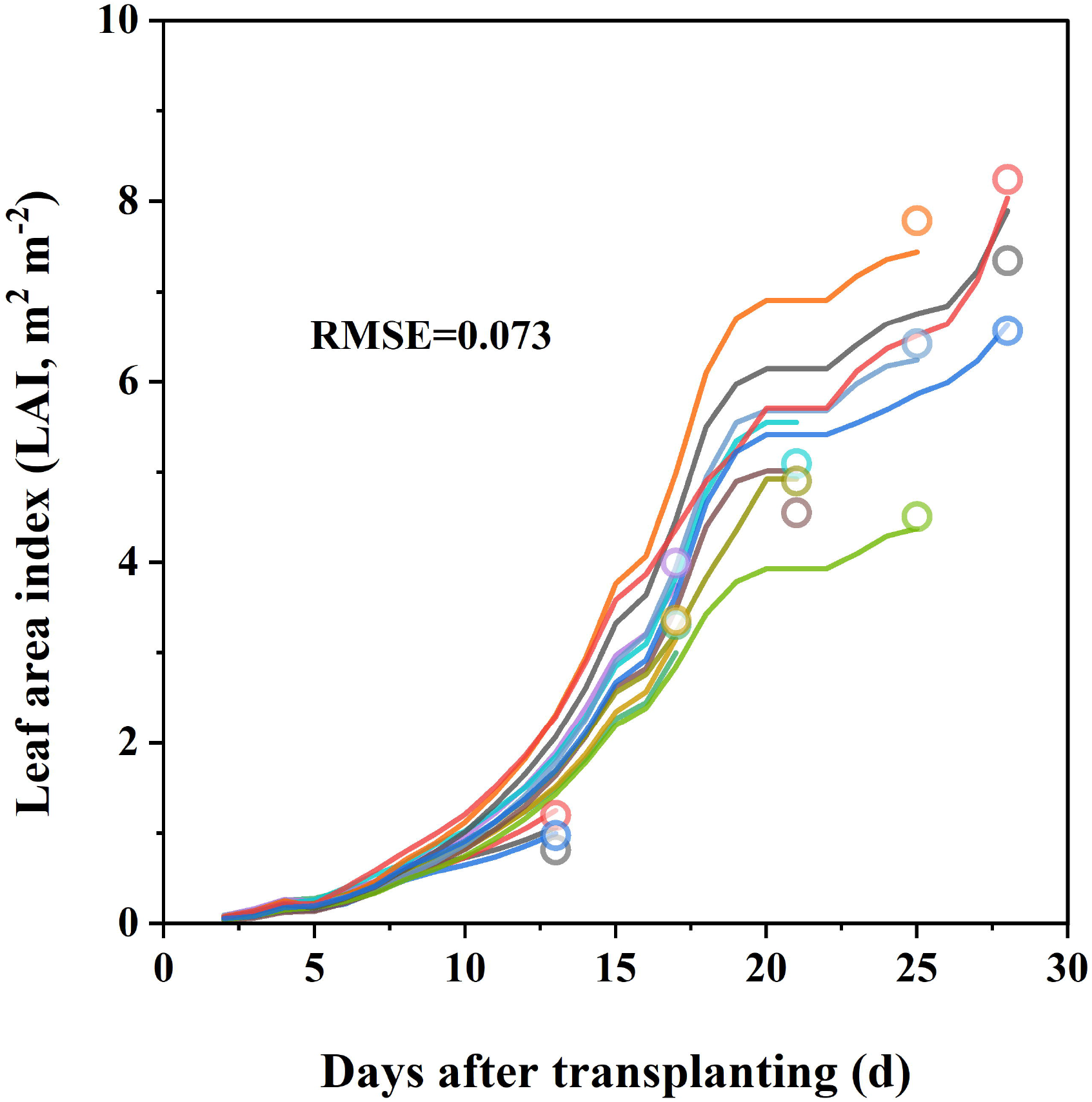
Comparison of estimated and measured leaf area index (LAI). LAI was estimated by the methods described in the “Leaf area index (LAI) estimation” section of SI (shown as lines). At 13,17, 21, 25 and 28 days after transplanting, leaf tissues from 3 experiment units of Plantarray were destructively removed from watermelon plants, respectively. Leaf areas were measure by LA-S leaf area analyzer system (shown as circles). RMSE represents root mean square error.

## 3. Results and discussion

### 3.1. Coupled FPP and simulation models to estimate dynamic and momentary WUE

Our idea was to extend Sinclair and colleagues’ model (Eq. 1), a theoretical expression of WUE (Sinclair *et al*., 1984; Sinclair & Ghanem, 2020), to FPP data for dynamic WUE estimation (Eq. 2). Prior to doing so, we tested the reliability of this approach by using a dataset obtained from wheat via the traditional low-throughput canopy gas exchange method (Song *et al*., 2016). During a 5-day period, canopy photosynthesis rate was found to peak much earlier than canopy transpiration rate before noon (Fig. S2a), which led to a higher WUE during the whole forenoon (square in Fig. S2b). The WUE_e_ estimated from the canopy transpiration rate and VPD at half-hourly intervals could well reproduce the measured WUE during the daytime (line in Fig. S2). At the daily step, day-to-day variations of both measured and estimated WUE were observed (triangles in Fig. S2), which were highly consistent as indicated by the tiny root mean square error (RMSE) of 0.00137 g g^-1^. These results proved that the chosen WUE model was suitable for dynamic WUE estimation.

In our high-throughput FPP data, the daily estimated WUE (WUE_e_, g DW/g H_2_O) calculated by Eq.2 based on the Tr and VPD continuously acquired by the Plantarray system, was found to fluctuate across the course of the experiment and varied significantly by genotype (Fig. 3a, similar result during spring growth season was shown in Fig. S3). During the WI phase, WUE (g FW/g H_2_O) within this period could be measured based on traditional method, i.e., accumulated daily transpiration dividing by accumulated biomass gain. The measured WUE based on fresh weight returned similar result of genetic difference to estimated WUE_e_ based on dry weight (Fig. 3b). Among the three genotypes, “HA” possessed the highest WUE across all phases, in particular late WD stage, followed by “JC5” and “PI296341-FR” (Fig. 3a, Fig. S3), in accord with the empirical knowledge that wild varieties tend to possess a higher level of drought resistance by sacrificing more biomass production (Kawasaki *et al*., 2000).

**Figure 3.**
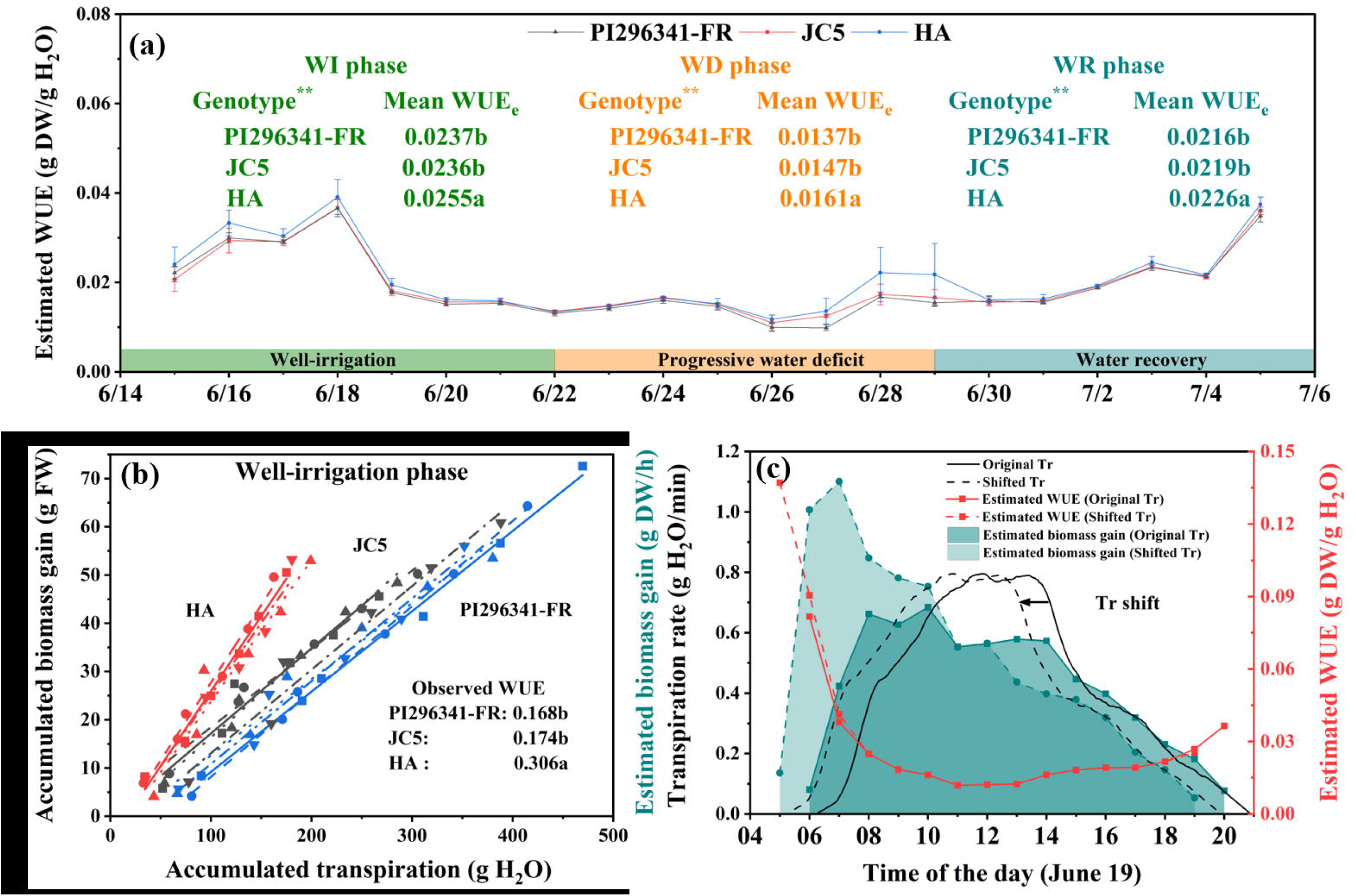
Measured and estimated water use efficiency (WUE), and estimated biomass gain based on models. a, daily estimated WUE (WUEe, g DW g-1 H2O) using mechanism models based on continuously recorded transpiration rate and VPD by Plantarray during the summer-season experiment in 2021. Multiple comparison results among three genotypes were shown as mean values with lowercase letters to show statistically significant differences for the well-irrigation (WI), progressive water deficit (WD) and water recovery (WR) phases, respectively. b, measured WUE (g FW g-1 H2O) based on accumulated biomass and transpiration during the summer-season experiment in 2021. Accumulated biomass (g FW) and transpiration (g H2O) were calculated according to daily biomass gain and daily transpiration measured by “Plantarray” system during WI phase. Accumulated biomass of PI296341-FR, JC5 and HA were fitted to a linear function of accumulated transpiration for individual experiment units, and multiple comparison results among three genotypes were shown. c, dynamic Tr (every 3 minutes), hourly WUE and biomass gain estimated based on models, taking the data from an experiment unit in June 19 for instance. A left-ward shift of dynamic Tr was applied to artificially advance it for 1 h. Corresponding WUE and biomass gain were estimated and shown as dotted lines.

When shortening the interval i in Eq. 2, the higher-resolution WUE (in the extreme case, the instantaneous or momentary WUE) and its diurnal variation could be estimated. In Fig. 3c (red solid line), we show that the hourly WUE_e_ calculated from measured Tr and VPD peaked in the early morning (6:00 am) and decreased until reaching a stable level at noon, and then rebounded slightly in the afternoon, which was consistent with the gas-exchange measurements elsewhere at the single leaf scale (Peddinti *et al*., 2019; Yang *et al*., 2019) or at the canopy scale.

### 3.2. Inferred sensitivity of Tr to Rad/VPD through inverse modeling

With the diurnal data of Tr and VPD, we were also able to conduct a simulation analysis by artificially advancing the dynamic Tr for 1 hour (black solid line shift to black dotted line) across a whole day (Fig. 3c), which revealed an increase of daily WUE_e_ and Biomass_e_ by 29.9 % and 20.0 %, respectively. The hourly WUE_e_ (red line) and Biomass_e_ (blue shadows) indicated that enhancing WUE during the early morning contributed greatly to the biomass gain (Fig. 3c). According to Eq. 1 and Eq. 2, the interpretation of this result is straightforward: without sacrificing daily accumulative Tr, more Tr in the low-VPD morning time decreases the integral value of daily VPD weighted by Tr, thus contributing to the increase of whole-day WUE and biomass gain. Such relatively small, daily improvement will accumulatively lead to an improved seasonal yield. Therefore, the pattern of diurnal Tr, even under the same daily cumulative transpiration conditions, is considered to reverse daily WUE.

To disentangle the intrinsic traits related to sensitivity of Tr to major environmental factors that were hidden within the daily Tr profiles, the modified Penman–Monteith model (Eq. 4) was employed to fit the dynamic Tr with the measured Rad and VPD, as well as the estimated daily LAI. LAI is often estimated as a function of thermal time, i.e. growing degree days (Medrano *et al*., 2005; Jo & Shin, 2021); however, such approch fell short of the need in our study, as LAI of individual plants differed remarkably which would influence the parameterization of Eq. 4. Here, daily transpiration of individual plants monitored by Plantarray in a non-destructive way was used to calibrate LAI in Eq. 4, a strategy that has been widely applied in coupling remote sensing and crop modeling (Zhao et al., 2013). Meanwhile, we measured the dynamic LAI of watermelon plants destructively during canopy development. As shown in Fig. 2, the estimated and measured LAI at each sampling time were similar (RMSE= 0.073), corroborating the robustness of our methods. Following that, two genotype-specific parameters in this model, *A* and *B*, which were essentially the S_Tr-Rad_ and S_Tr-VPD_, were estimated from curve fitting of transpiration rate with the input of measured Rad, VPD and estimated LAI, respectively. The fitted transpiration dynamics well reproduced the measured diurnal transpirations, as evidenced by the high correlation (R^2^=0.94) between them and the low total RMSE of only 0.05 g/min, taking the commercial cultivar “JC5” for example (Fig. 4a). S_Tr-Rad_ and S_Tr-VPD_ were thus inferred via inverse modeling in a daily step as shown in Fig 4b & c (similar result during spring growth season was shown in Fig. S4). We conducted two-way ANOVA for the daily S_Tr-Rad_ and S_Tr-VPD_ within each phase of WI, WD and WR, and the result indicated genotypic variation and dramatic change in each of them across the treatment phases (*P* < 0.01, Fig. 4b, c, Fig. S4). In general, S_Tr-Rad_ decreased as the water deficit progressed and rapidly recovered when water was resumed (Fig. 4b, Fig. S4a); S_Tr-VPD_, however, decreased quickly on the initial days of water deficit, and continued to decrease gradually on the following days until reaching a rather stable level even when irrigation was resumed (Fig. 4c, Fig. S4b). Among the three genotypes, “HA” exhibited high S_Tr-Rad_ only at the WD phase and remarkedly low S_Tr-VPD_ particularly at the WI stage in comparison to the remaining two genotypes (Fig. 4b&c, Fig. S4), indicating that plasticity of S_Tr-Rad_ and S_Tr-VPD_ in response to the environment is important in shaping the genotypic-specific dynamic WUE patterns.

**Figure 4.**
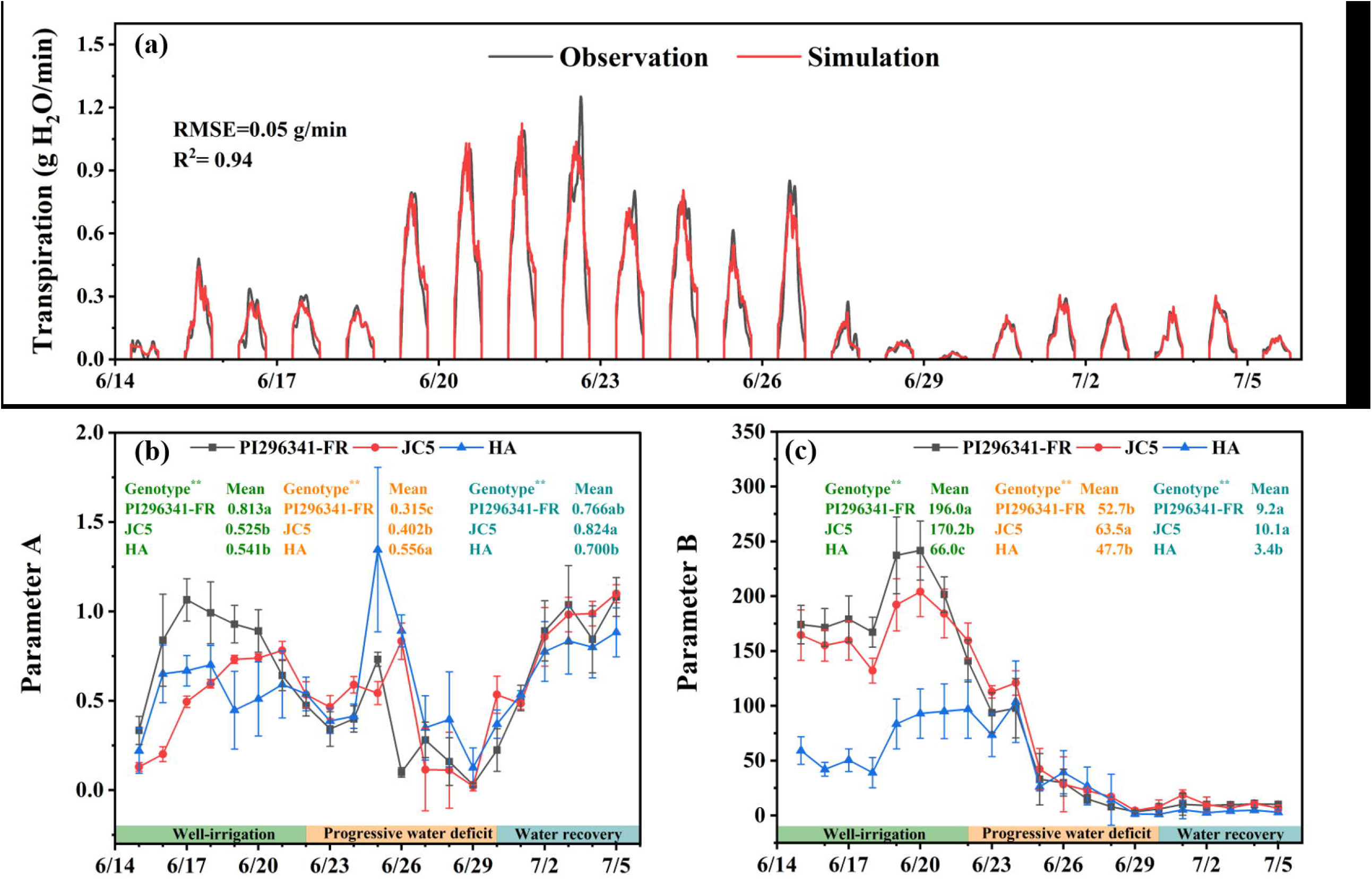
Simulation of diurnal transpiration by using the modified Penman-Monteith model. a, observed and simulated diurnal variation of transpiration rate during summer-season experiment in 2021, taking an experiment unit of JC5 for instance. The observations and simulations were both 3-min steps. b-c, fitted daily parameter A or S_Tr-Rad_ (b) and parameter B or S_Tr-VPD_ (c) by modified Penman-Monteith model during summer-season experiment in 2021. Multiple comparison results among three genotypes (PI296341-FR, JC5, and HA) were shown as mean values with lowercase letters to show statistically significant differences for the well-irrigation (WI), progressive water deficit (WD) and water recovery (WR) phases, respectively.

### 3.3. Transpiration ideotype design based on S_Tr-Rad_ and S_Tr-VPD_

Based on the results above, we propose that an optimized combination of S_Tr-Rad_ and S_Tr-VPD_ is important for balanced Tr, WUE, and biomass gain. Simulation models are powerful tools to support breeding because of their capability to reveal the interactions between each component within a system. Therefore, we conducted simulations with the inputs of 5-day meteorological data from each of the WI and WD phases (Fig. 5a, c) and the artificial combinations of S_Tr-Rad_ and S_Tr-VPD_ extremes (i.e., highest parameter A × lowest parameter B, lowest A × lowest B, highest A × highest B, and lowest A × highest B in the model) adopted from the 3 watermelon genotypes averaged over each 5-day course (Fig. 5b, d), where LAI was fixed to 3 to maintain a fair canopy size. Since Rad peaked much earlier than VPD in the morning (Fig. 5a, c), high A and low B resulted in high Tr during the low VPD intervals within a day and consequently a low integrated value of daily VPD weighted by Tr (Eq. 1), leading to high WUE (red line in Fig. 5b, d).

**Figure 5.**
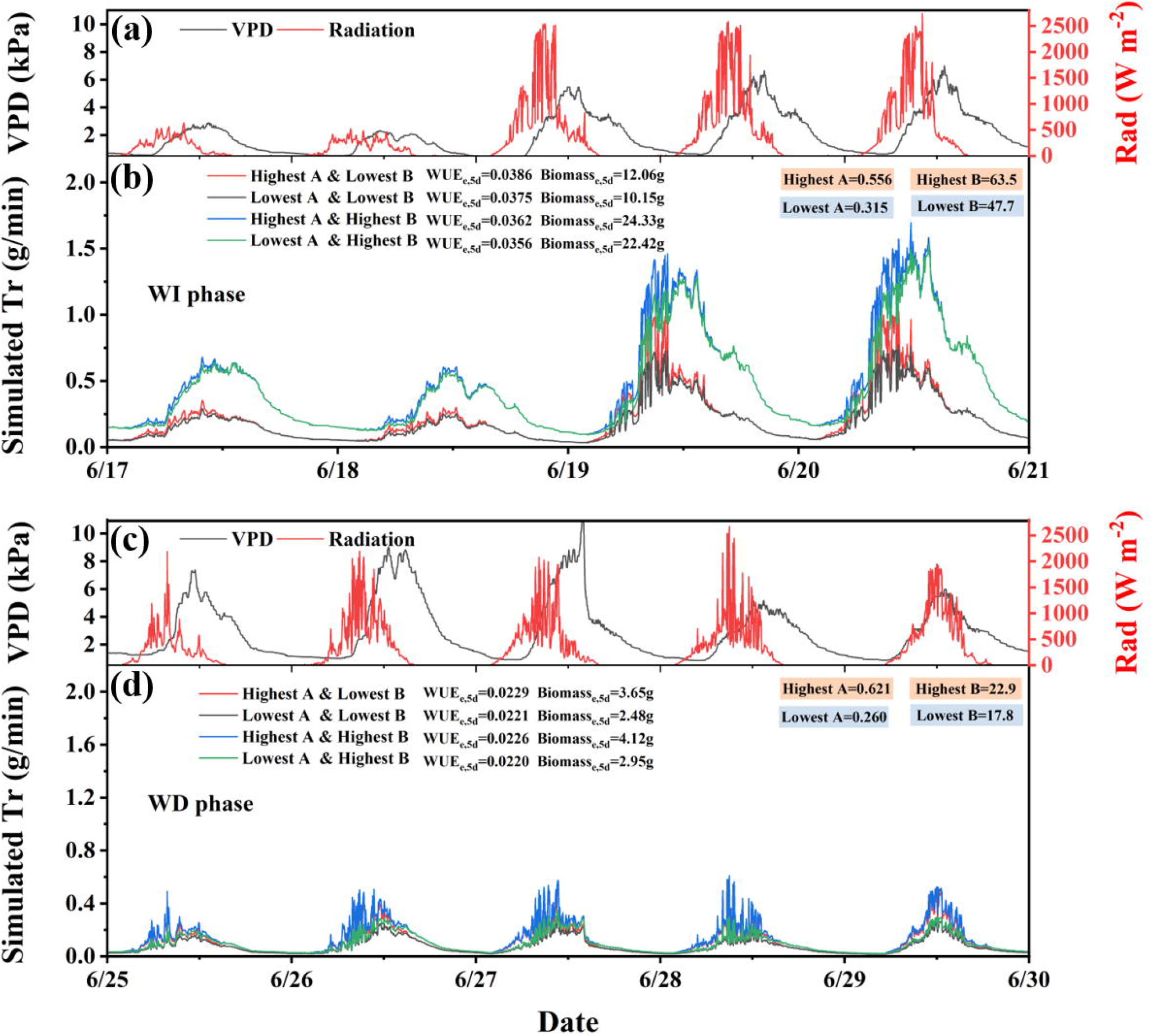
Diurnal variation of Rad and VPD, and the responsive patterns of simulated Tr to different combinations of parameter A and B. a, c five-day Rad and VPD (normalized to 0-1 scale) recorded at 3-min intervals, which were selected from the well-irrigation (WI, 6.17-6.21, a) and progressive water deficit (WD, 6.25-6.29, c) phase, respectively. The LAI was set to a constant of 3 for all simulations. Mean values of parameters A and B for different genotypes (PI296341-FR, JC5, and HA) were calculated over each 5-day course, and the maxima and minima formed the four combinations as shown in the graph. Mean estimated WUE and biomass gain over 5-day course were denoted as WUE_e,5d_ and Biomass_e,5d_, respectively.

Further, the estimated biomass gains of different combinations were shown in Fig 5b, d. During the WI phase, highest A × highest B and lowest A × highest B both resulted in much larger Tr, which produced approximately double the biomass gain compared to the remaining combinations despite their lower WUE (Fig. 5b). While during the WD phase, the highest A × highest B still obtained the highest biomass gain benefitting from the highest Tr (blue line in Fig. 5d), but the highest A × lowest B obtained a second highest biomass gain. Therefore, by considering both WUE and Tr, we put forward the following principles for transpiration ideotype design: in the well-irrigated agricultural areas, crop genotypes with high S_Tr-Rad_ and high S_Tr-VPD_ is favorable by producing superior yield; in the arid agricultural areas where water-saving is a necessity, genotypes with high S_Tr-Rad_ and low S_Tr-VPD_ is desired for its marvelously reduced transpiration yet still considerable yield; in the intermittent drought-prone areas where irrigation and water shortage alternate, genotypes showing strong plasticity in S_Tr-Rad_ and S_Tr-VPD_ that are suited to the optimized balance of yield and water consumption will be desired.

## 4. Conclusions

We demonstrate that daily to hourly or even instantaneous WUE can be estimated by coupling existing WUE models with the dynamically recorded transpiration and VPD from high-throughput FPP, which breaks the bottleneck of dynamic and momentary WUE acquisition. We further demonstrate that daily WUE is affected by the diurnal transpiration pattern of a plant, which is tightly linked to the genotype-dependent sensitivity of Tr to Rad and VPD. A quantitative approach was developed to infer these transpiration sensitivity parameters. We propose that it is beneficial to increase WUE by lowering S_Tr-VPD_ while offsetting its negative impact on Tr by increasing S_Tr-Rad_. Although in previous studies high-WUE crops selected by carbon isotope discrimination generally failed to match the demand of luxuriant growth (Blum, 2005; Blum, 2009), the high-WUE trait is not necessarily linked to growth limitation (Marguerit *et al*. (2014), which lays the foundation for synergistic improvement of WUE and yield. FPP-enabled phenomic selection will help catch the desired elite individuals from enormous germplasm lines, e.g. those with substantially high S_Tr-Rad_ under both well-watered and water scarcity conditions.

## Acknowledgments

We thank the supports of National Key Research & Development Program of China (2021YFE19800), the Research & Development Program of Ningbo (2021Z006) and the Key Scientific and Technological Grant of Zhejiang for Breeding New Agricultural Varieties (2021C02066-5, 2021C02067-7).

