## Supplementary material for "Coupled functional physiological phenotyping and simulation model to estimate dynamic water use efficiency and infer transpiration sensitivity traits": Fig. S1

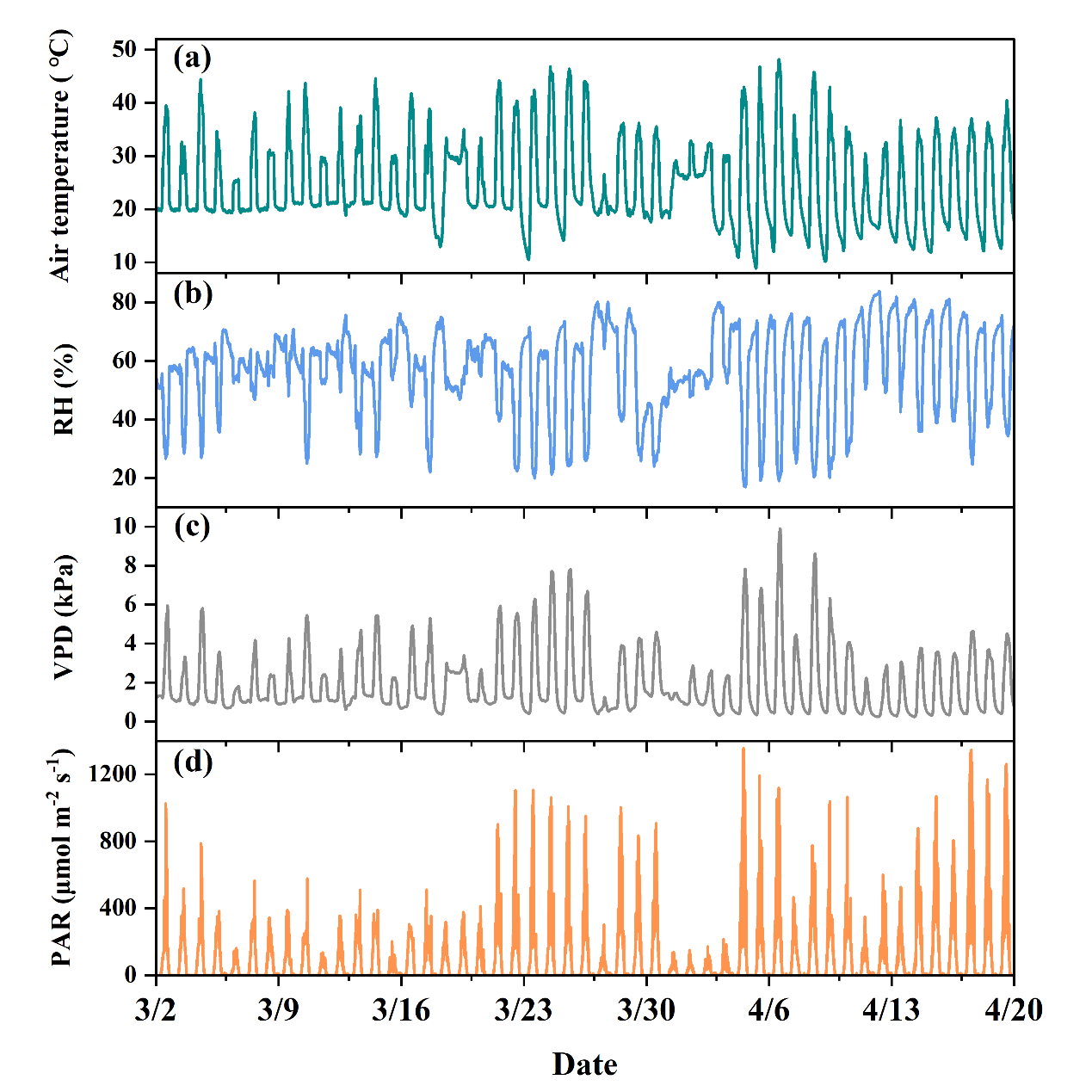
**Figure S1** Parameters of the soil-plant-atmosphere continuum monitored by the high-throughput phenotyping platform “Plantarray”. Meteorological parameters, i.e., air temperature (T_air_, ℃, a), relative humidity (RH, %, b), vapor pressure difference (VPD, kPa, c), and photosynthetic active radiation (PAR, μmol m^-2^ s^-1^, d) were obtained during the spring-season experiment in 2021.
