## Supplementary material for "Coupled functional physiological phenotyping and simulation model to estimate dynamic water use efficiency and infer transpiration sensitivity traits": Fig. S2

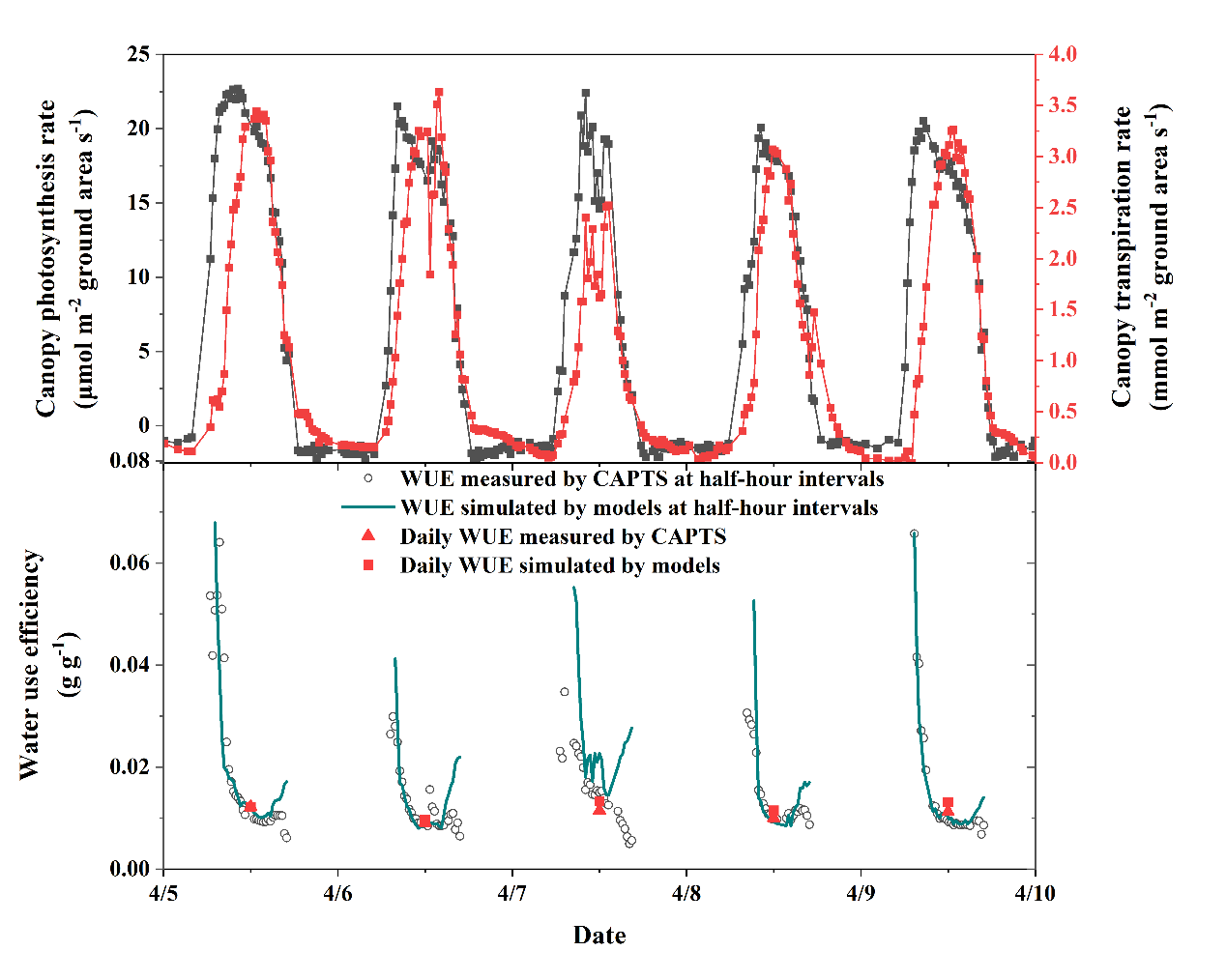

**Figure S2** WUE calculated by measured canopy gas change parameters and estimated by simulation models. a. Canopy photosynthesis rate (left vertical axis) and transpiration rate (right vertical axis) measured by canopy photosynthesis and transpiration measurement system (CAPTS). b. Canopy WUE calculated from measured canopy photosynthesis rate and transpiration rate at half-hour intervals (circle point) and daily intervals (triangle), and simulated by Eq. 2 with measured vapor pressure deficit (VPD) and canopy transpiration rate at half-hour intervals (line) and at daily intervals (square). The datasets were obtained from wheat in the field in the year of 2021. *K*_d_ in Eq. 2 was estimated as 3.5KPa.
