## Supplementary material for "Coupled functional physiological phenotyping and simulation model to estimate dynamic water use efficiency and infer transpiration sensitivity traits": Fig. S3

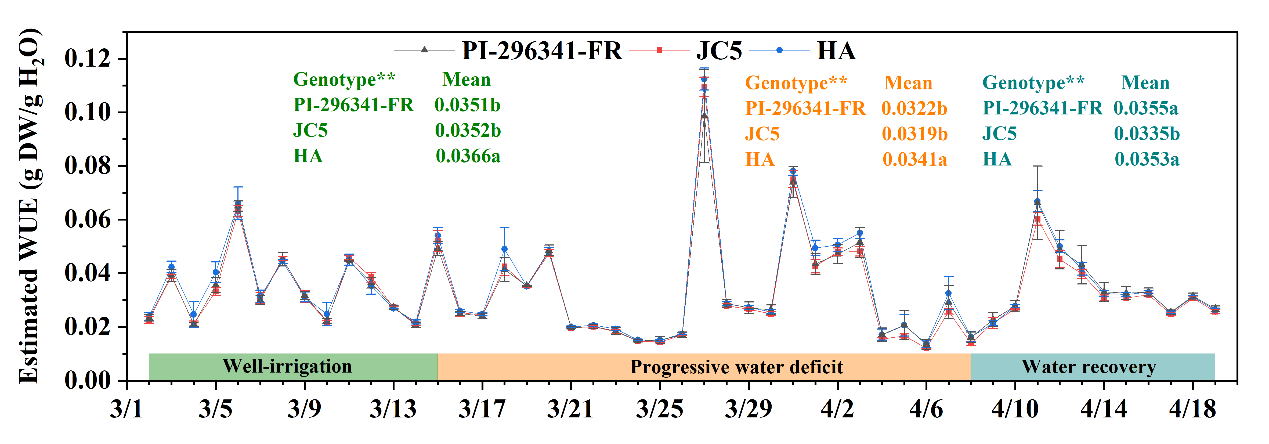

**Figure S3** Estimated water use efficiency (WUE_e_). Daily estimated WUE (WUE_e_, g DW g^-1^ H_2_O) was calculated based on the Eq. 2 with the observed transpiration and VPD data from the spring-season experiment in 2021. Multiple comparison results among three genotypes (WT, JC5, and HA) were shown as mean values with lowercase letters to show statistically significant differences for the well-irrigation (WI), progressive water deficit (WD) and water recovery (WR) phases, respectively.
