## Supplementary material for "Coupled functional physiological phenotyping and simulation model to estimate dynamic water use efficiency and infer transpiration sensitivity traits": Fig. S4

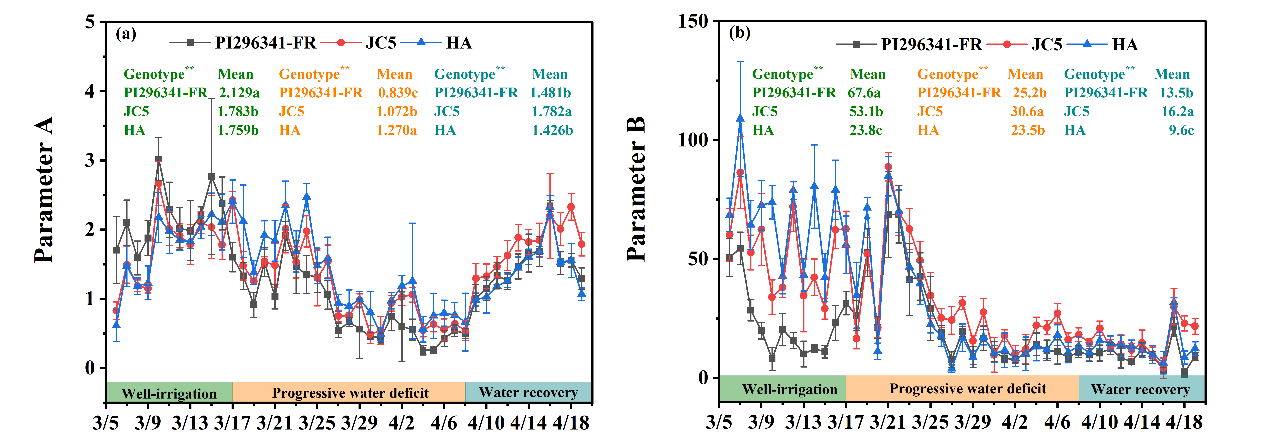

**Figure S4** Fitted daily parameter A or S_Tr-Rad_ (a) and parameter B or S_Tr-VPD_ (b) by modified Penman-Monteith model during spring-season experiment in 2021. Multiple comparison results among three genotypes (PI296341-FR, JC5, and HA) were shown as mean values with lowercase letters to show statistically significant differences for the well-irrigation (WI), progressive water deficit (WD) and water recovery (WR) phases, respectively.
